# ADS-J1 Disaggregates Semen-derived Amyloid Fibrils

**DOI:** 10.1101/292342

**Authors:** J. Li, Z. Yang, H. Liu, Y. Lan, T. Zhang, W. Li, T. Qi, Y. Qiu, L. Li, X Zhou, S. Liu, S. Tan

**Keywords:** HIV-1, SEVI, amyloid fibril, disaggregate, molecular dynamic simulation

## Abstract

Semen-derived amyloid fibrils, composing SEVI (semen-derived enhancer of viral infection) fibrils and SEM1 fibrils, could remarkably enhance HIV-1 sexual transmission and thus, are potential targets for the development of an effective microbicide. Previously, we found that ADS-J1, apart from being an HIV-1 entry inhibitor, could also potently inhibit seminal amyloid fibrillization and block fibril-mediated enhancement of viral infection. However, the remodeling effects of ADS-J1 on mature seminal fibrils were unexplored. Herein, we investigated the capacity of ADS-J1 to disassemble seminal fibrils and the potential mode of action by applying several biophysical and biochemical measurements, combined with molecular dynamic (MD) simulations. We found that ADS-J1 effectively remodeled SEVI, SEM1_86-107_ fibrils and endogenous seminal fibrils. Unlike epi-gallocatechin gallate (EGCG), a universal amyloid fibril breaker, ADS-J1 disaggregated SEVI fibrils into monomeric peptides, which was independent of oxidation reaction. MD simulations revealed that ADS-J1 displayed strong binding potency to the full-length PAP_248-286_ via electrostatic interactions, hydrophobic interactions and hydrogen bonds. ADS-J1 might initially bind to the fibrillar surface and then occupy the amyloid core, which eventually lead to fibril disassembly. Furthermore, the binding of ADS-J1 with PAP_248-286_ might induce conformational changes of PAP_248-286_. Disassembled PAP_248-286_ might not be favor to re-aggregate into fibrils. ADS-J1 also exerts abilities to remodel a panel of amyloid fibrils, including Aβ_1-42_, hIAPP_1-37_ and EP2 fibrils. ADS-J1 displays promising potential to be a combination microbicide and an effective lead-product to treat amyloidogenic diseases.

## INTRODUCTION

Sexual transmission, either via vaginal intercourse or via anal intercourse, is the predominant route of human immunodeficiency virus type I (HIV-1) acquisition. Both oral and topical pre-exposure prophylaxis (PrEP) are promising approaches to reduce HIV-1 sexual transmission (1). Although oral administration of tenofovir disoproxil fumarate/emtricitabine regimen has been a significant progress in HIV-1 prevention (2), the use of topical PrEP may be preferential over oral PrEP due to the rapid onset of protective drug levels at mucosal sites following administration and the potential reduction of systemic adverse effects (3). Development of topical microbicide is still urgent and challenging.

Semen, as an important carrier of HIV-1, plays critical roles in HIV-1 sexual transmission worldwide (4). Semen contains endogenous amyloid fibrils that could potently enhance HIV-1 infectivity and impair the anti-HIV activities of antiretroviral drugs (5, 6). The seminal amyloid fibrils compose of proteolytic peptide fragments of prostatic acid phosphatase (PAP), a protein produced by the prostate gland. The best characterized seminal amyloid is formed by PAP_248-286_ and termed SEVI (semen-derived enhancer of virus infection) (7). Later, a second serial of seminal fibrils were identified from prostate specific antigen (PSA)-generated fragments of semenogelin (SEM1 and SEM2) and PAP_85-120_ (8, 9) which could also facilitate HIV-1 infection. These amyloid fibrils contain a high abundance of positively charged arginine and lysine residues, which help to promote virion attachment and subsequent infection by allowing the virus to overcome the electrostatic repulsion between the negatively charged viral and cellular membranes (10).

Microbicide, simultaneously targeting virions and seminal amyloid fibrils, might be an innovative strategy to slow the global spread of HIV-1 (11, 12). Previously, our study showed that ADS-J1, an HIV-1 entry inhibitor targeting gp120 and gp41 (13, 14), displayed potential abilities to counteract seminal amyloid fibril-mediated enhancement of viral infection. Due to its sulfonic groups, ADS-J1 binds to peptide PAP_248-286_ and inhibits SEVI fibril formation. In addition, it coats on and shields the positive charges of mature SEVI fibrils, thus antagonizing fibril-mediated enhancement of HIV-1 infection (3). Unlike polyanions, which have failed in clinical trials, ADS-J1 does not accelerate SEVI fibril formation at low concentrations (15). Importantly, combination of ADS-J1 with antiretroviral (ARV)-based candidate microbicides displays synergistic and additive effects against HIV-1 infection in semen, with little cytotoxicity to vaginal epithelial cells. With the anti-amyloid and anti-viral pharmaceutical effects, our results were in support of the potential application of ADS-J1 as a combination microbicide.

However, the remodeling effects of ADS-J1 on mature SEVI fibrils have been elusive. Endogenous mature fibrils are already formed and abundant in semen (6). Thus, inhibition of the initial fibril formation would seem impractical or even futile during sexual intercourse. Although ADS-J1 could coat the amyloid surface and prevent interactions with the virus and cell surface, fibrils might also disassociate with ADS-J1 and gain chances to facilitate viral infection. A more desirable feature of ADS-J1 that could rapidly deconstruct seminal fibrils and thereby disrupt structures that promote HIV-1 infection is highly warranted. In this study, we intended to investigate the potential effects of ADS-J1 on remodeling seminal mature fibrils and study the underlying mechanism of action.

## MATERIALS AND METHODS

### Materials

Thioflavin T (ThT), EGCG and polybrene were purchased from Sigma (St, Louis, MO). Superoxide dismutase (SOD) was purchased from Beyotime Biotechnology (Shanghai, China). The Proteostat amyloid plaque detection kit was purchased from Enzo Life Sciences (Plamouth Meeting, PA). Peptides, including PAP_248-286_, SEM1_86-107_, hIAPP_1-37_ and EP2 peptide (> 95% purity), were all synthesized by Scilight-Peptide (Beijing, China). Polyclonal rabbit antibody against PAP_248-286_ was produced by AbMax Biotechnology Co, Ltd. (Beijing, China). Semen (SE) samples were obtained from healthy members of laboratory with written informed consent. The protocol was approved by Human Ethics Committee of Southern Medical University, China. Seminal fluid (SE-F), represent the cell-free supernatant of SE, was collected by centrifugation for 30 min at 10,000 rpm at 4°C and stored in 1 ml aliquots at −20°C. TZM-bl cells were obtained from the Nation Institutes of Health AIDS Research and Reference Reagent Program. CCR5-tropic SF162, CXCR4-tropic NL4-3 and dual tropic infectious clone plasmids were kindly provided by Jan Münch of Ulm University, Ulm, Germany. Viruses were prepared as previously described (15). Briefly, viruses were produced by polyethyleneimine (PEI) mediated transfection of 293TX cells with DNA proviral expression plasmids. Viruses were collected 48 h later. ADS-J1 was purified by high-performance liquid chromatography (HPLC) from the crude material provided by Ciba Specialty Chemical Corp. (High Point, NC).

### Fibrils preparation

Lyophilized peptides of PAP_248-286_ and SEM1_86-107_ were dissolved at a concentration of 2200 μM in phosphate-buffered saline (PBS) and stored at −20°C. Fibrils formation was initiated by agitated at 37°C for 1 to 3 days at 1,400 rpm with an Eppendorf Thermomixer. Fibrils of Aβ_1-42_ and hIAPP_1-37_ were prepared as previously described (16). EP2 lyophilized peptides were dissolved in dimethyl sulfoxide (DMSO) and stored at −20°C. The peptides were dissolved in PBS and fibril formed immediately.

### ThT staining assay

Preformed SEVI, SEM1_86-107_, Aβ_1-42_, hIAPP_1-37_ and EP2 fibrils were agitated with or without ADS-J1 at 1,400 rpm at 37°C with an Eppendorf Thermomixer. At the indicated time points, 10 μl aliquots were withdrawn from each sample for ThT staining as previously reported (15). Briefly, 10 μl aliquots samples were mixed with 90 μl ThT (50 μM) in 96-well black plate and the fluorescence (excitation at 440 nm, emission at 482 nm) was measured immediately with a fluorescence plate reader (Infinite M1000, Tecan). To assess the effects of ADS-J1 on disaggregating endogenous seminal fibrils, ADS-J1 at different concentrations or PBS was agitated with whole SE-F, and samples were withdrawn at different time points to detect fibril integrity by ThT staining as described above.

### Transmission electron microscopy (TEM)

Peptide samples were incubated in the absence or presence of ADS-J1 as described for the ThT fluorescence measurements. Samples were removed at different time points and absorbed onto glow-discharge carbon coated grids for 2 min. Excess samples were removed and the grids immediately were stained with 2% phosphotungstic acid for 2 min. The fibrils were visualized with an H-7650 transmission electron microscopy (Hitachi Limited, Tokyo, Japan).

### Analysis of tricine-SDS-PAGE

Mature SEVI fibrils were agitated with ADS-J1 or EGCG at 37°C at 1,400 rpm for 12 h. Samples were centrifuged at 12,000 rpm for 15 min. The remodeling products in the supernatant were analyzed on 15% Tricine SDS PAGE gels. Gels were stained with Coomassie blue.

### HIV-1 infection assays

TZM-bl cells (1×10^3^) in 100 μl cell culture medium were seeded in 96-well flat-bottom plates the day before infection. Mature fibrils or whole SE-F were agitated in the presence or absence of ADS-J1 at various concentrations. At different time points, 60 μl samples were collected and centrifuged at 12,000 rpm for 10 min to remove the remaining free ADS-J1. The pellets were dissolved in 60 μl serum-free culture medium and mixed with 60 μl viruses (2 ng of p24 antigen). After 10 min, 100 μl of the mixtures were added to TZM-bl cells. After a 3-h incubation period, unbound viruses were removed and cells were replenished with fresh medium. Luciferase activity was measured 48-h post-infection. SEVI or SEM1_86-107_ fibrils were tested at a final concentration of 11 μM, while SE-F was used at a final dilution of 1:100.

### Measurement of Zeta potential and particle size

Fibrils were agitated with 2 or 4-fold excess of ADS-J1 at 1,400 rpm at 37°C for 4 h. Fibrils were then centrifuged at 12,000 rpm for 10 min. The pellets were resuspended in 1 mM KCl. Both zeta potential and particle size were measured using the Zeta Nanosizer (Malvern, Worcestershire, UK).

### Competition experiments

ADS-J1 (110 μM) was incubated with polybrene at different concentration (0 μM, 110 μM, 220 μM, 440 μM) at 37°C for 30 min and then the mixtures were centrifuged for 15 min at 12,000 rpm. The supernatants were agitated with SEVI. Fibrillar signal was examined by ThT staining assay as mentioned above.

### SOD experiments

SEVI was agitated with ADS-J1 or EGCG in the presence or absence of SOD at 37 °C for 4 h. The ability of ADS-J1 or EGCG to remodel SEVI fibrils in the presence of SOD was evaluated by ThT staining method.

### MD simulation

MD was used to investigate the interaction between PAP_248-286_ and ADS-J1. The original structure coordinates for PAP_248-286_ was retrieved from the Protein Data Bank (PDB ID: 2L3H) (17). The initial binding between PAP_248-286_ and ADS-J1 were searched by Autodock vina (18).

All of MD simulations were performed 100 ns using the GROMACS 4.5.3 software package (19) together with CHARMM27 force field (20). The system was placed in the center of a cubic box that extended 6 Å from the structure on each side. TIP3P water molecules were added together with sodium and chloride ions to offer isotonic condition and neutralize the whole system (21). Then, the system was energy minimized for 3000 steps by releasing all the constraints. All bond lengths were constrained by LINCS algorithm (22). The electrostatic interaction was calculated by the particle mesh Ewald (PME) method (23). The MD temperature was controlled at 300 K and pressure was set at 1 atm. The MD simulation trajectories were saved in every 2 ps for analysis.

The most representative structure and contact map in each simulation system was gained by cluster analysis with a cut-off value of 0.3 nm. The root-mean-square fluctuation (RMSF) value presents the fluctuations of C_α_ atoms. The structures from the simulation trajectories were visualized using YASARA (24).

## RESULTS

### Remodeling effects of ADS-J1 on SEVI and SEM1_86-107_ fibrils

To investigate the effects of ADS-J1 on remodeling preformed seminal amyloid fibrils, we first detected its ability to remodel SEVI and SEM1_86-107_ fibrils using ThT staining assay. In the absence of ADS-J1, 30 μM SEVI fibrils (Fig. 1 *A*) and SEM1_86-107_ fibrils (Fig. 1 *B*), which have been agitated in PBS at 37°C for 1 to 3 days at 1,400 rpm, remained constant binding with ThT, a specific dye of amyloid fibrils. It indicated both fibrils were stable in the buffer within the tested period. However, in the presence of ADS-J1, the ThT fluorescence intensity of fibrils decreased by around 50% in 4 h (Fig. 1, *A* and *B*). It suggested that the mature fibrils were converted into other form of species that lost the ability to bind ThT. Of note, ADS-J1 alone at tested concentrations did not influence ThT fluorescence (data not shown), indicating that ADS-J1 did not quench ThT fluorescence. Fibrillar disruption by ADS-J1 was also confirmed by TEM. Upon a 4-hour incubation, TEM images revealed that the control SEVI fibrils and SEM1 fibrils remained a dense of intact and branched fibrils. In contrast, treated with two fold excess of ADS-J1, SEVI fibrils were converted to smaller nonfibrillar species and SEM1 fibrils were converted to shorter fibrils (Fig. 1 *C*). ADS-J1 alone served as negative control. Particle size measurement was also applied to confirm our findings. The averaged particle sizes of SEVI and SEM1 fibrils were 3636 ± 208 nm and 2043 ± 307 nm respectively. Treated with 4 fold excess of ADS-J1, the fibrils were disaggregated into smaller pieces with averaged particle sizes of 506 ± 14 nm and 543 ± 6.9 nm respectively (Fig. 1 *D*).

**FIGURE 1.**
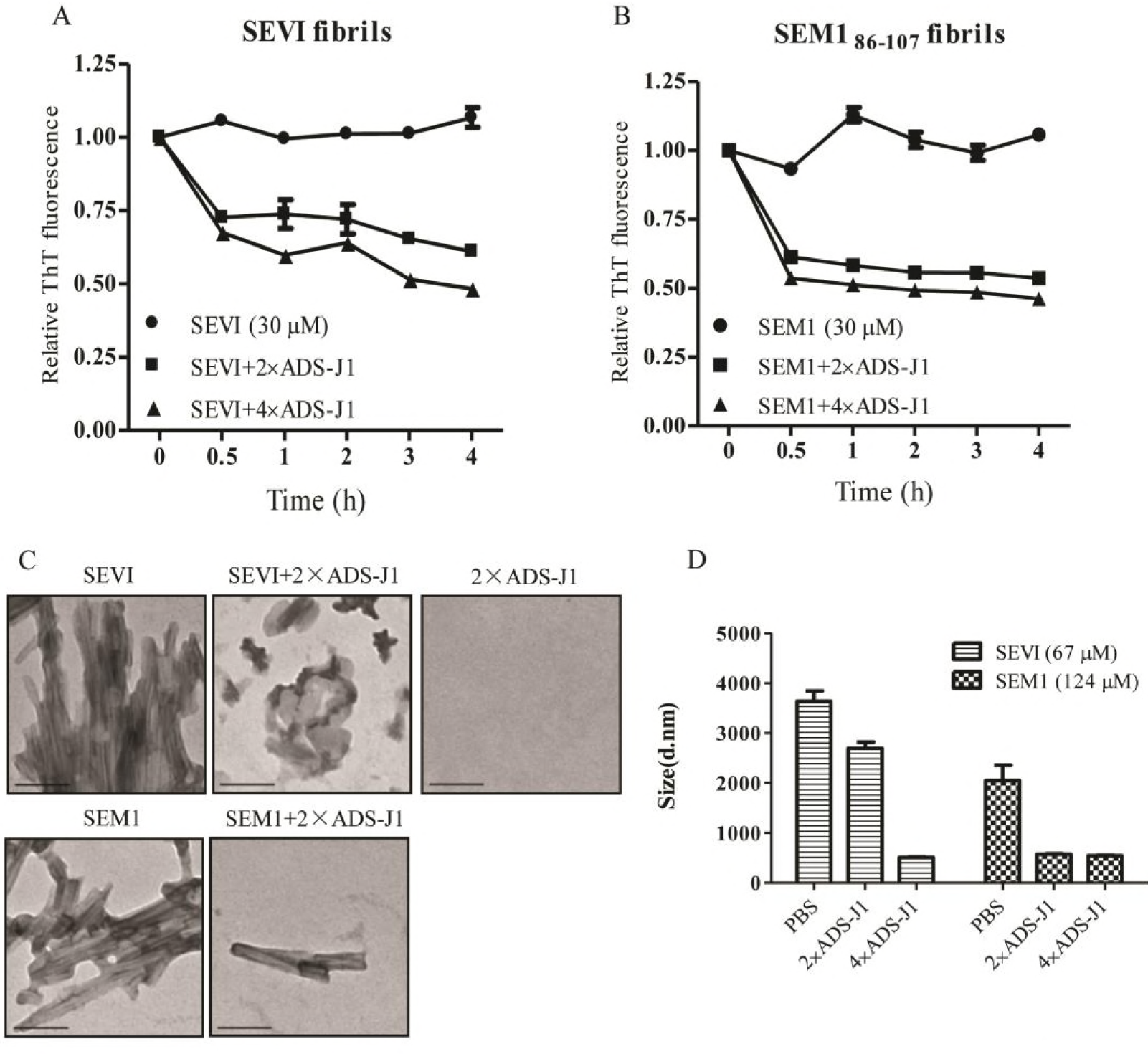
Remodeling effects of ADS-J1 on SEVI and SEM1_86-107_ fibrils. Fibrillar aggregates (30 μM) were exposed to different amounts of ADS-J1. Fibril integrities of SEVI (*A*) and SEM1_86-107_ (*B*) were detected by ThT staining at the indicated time points. Values are means ± SEM (n=3). (*C*) Preformed SEVI fibrils and SEM1_86-107_ fibrils (110 μM) were treated with a 2-fold excess of ADS-J1 or buffer for 4 h at 37 °C. Fibrillar shapes were monitored by TEM. Scale bar = 100 nm. (*D*) Particle sizes of SEVI and SEM1_86-107_ fibrils agitated in the presence or absence of ADS-J1 at 37 °C for 4 h were measured. Results shown are means ± SEM of triplicate measurements from one of the two independent experiments, which yielded equivalent results.

Viral infection experiments were conducted to determine the activity of the disaggregating products to enhance HIV-1 infection. We found that the remodeling products showed impaired ability to facilitate HIV-1 infection. Although different fibrils enhanced various viral infection to a different extent, SEVI and SEM1 fibrils exerted stable activity to enhance HIV-1 infection within 48 h (Fig. 2). ADS-J1 decreased the ability of SEVI (Fig. 2, *A*-*C*) and SEM1 fibrils (Fig. 2, *D*-*F*) to enhance HIV-1 infection in a dose and time-dependent manner. An 8-fold molar excess of ADS-J1 could completely diminished fibril-mediated enhancement of HIV-1 infection (Fig. 2). Taken together, our findings suggest that ADS-J1 effectively remodels SEVI and SEM1 fibrils and abrogates their infection enhancing property.

**FIGURE 2.**
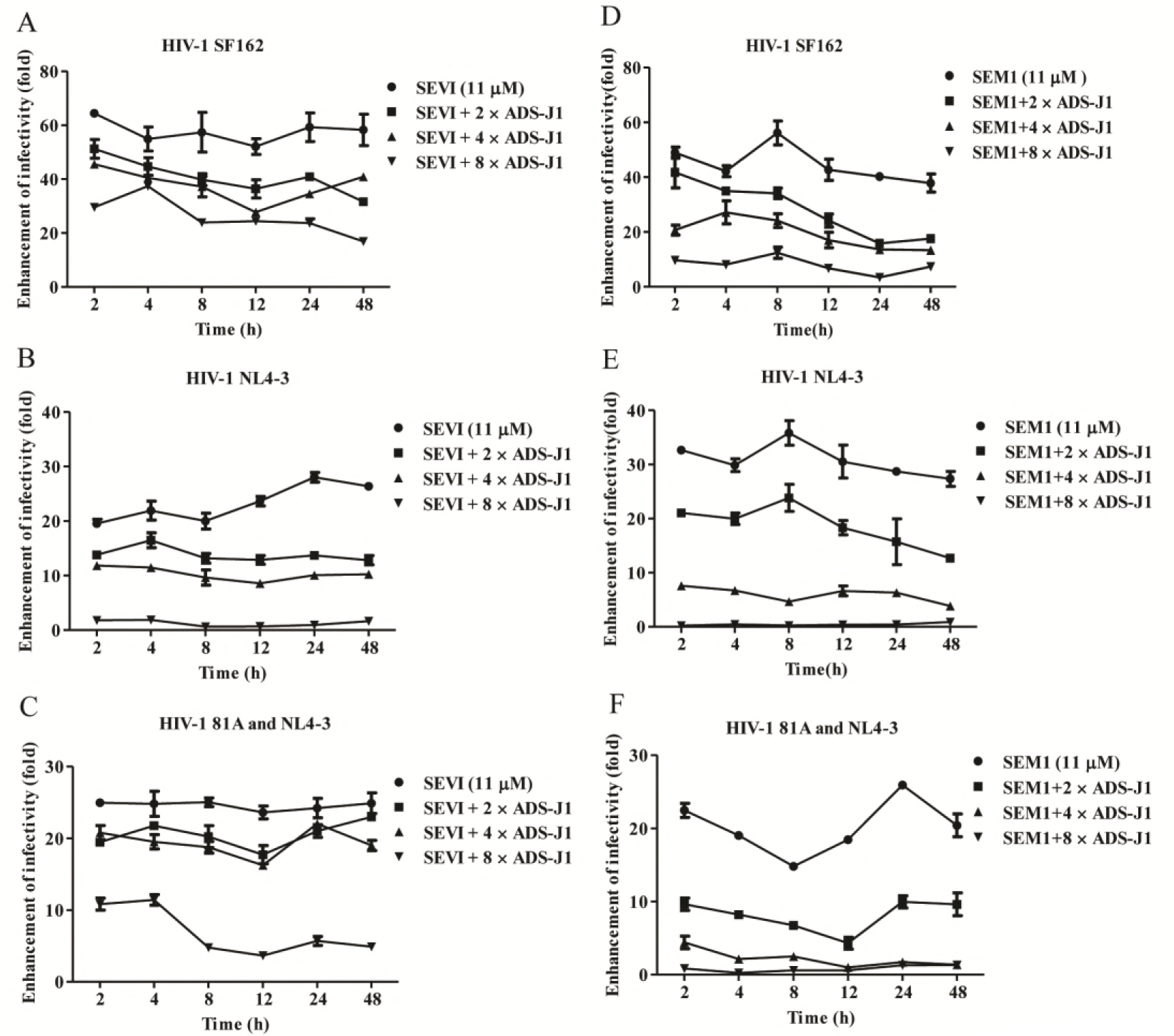
Abrogation of SEVI and SEM1 fibrils – mediated enhancement of HIV-1 infection by ADS-J1. SEVI (*A-C*) or SEM1_86-107_ fibrils (*D-F*) were agitated with increasing concentrations of ADS-J1. At the indicated time points, samples were collected and centrifuged. The pellets were mixed with CCR5-tropic HIV-1SF162 (*A* and *D*), CXCR4-tropic HIV-1 NL4-3 (*B* and *E*) or dual tropic HIV-1 (*C* and *F*). The rate of infection was monitored by measuring the luciferase activity in TZM-bl cells. The data are means ± SEM of three independent measurements.

### Remodeling effects of ADS-J1 on endogenous seminal fibrils

We then explored whether ADS-J1 could remodel endogenous fibrils in semen. We allowed fibrils to grow in semen by agitating SE-F for 8 h at 37°C, 1400 rpm, as evidenced by an increase in ThT fluorescence. Agitated SE-F was exposed to different concentrations of ADS-J1 for 4 h. ADS-J1 could dose-dependently decrease ThT fluorescent signal in semen (Fig. 3 *A*). Accordingly, after exposure to ADS-J1, SE-F gradually lost the ability to enhance HIV-1 infection (Fig. 3 *B*). ADS-J1 at 240 μM could completely block the enhancing property of SE-F (Fig. 3 *B*). These findings suggest that ADS-J1 could also remodel endogenous seminal amyloid fibrils.

**FIGURE 3.**
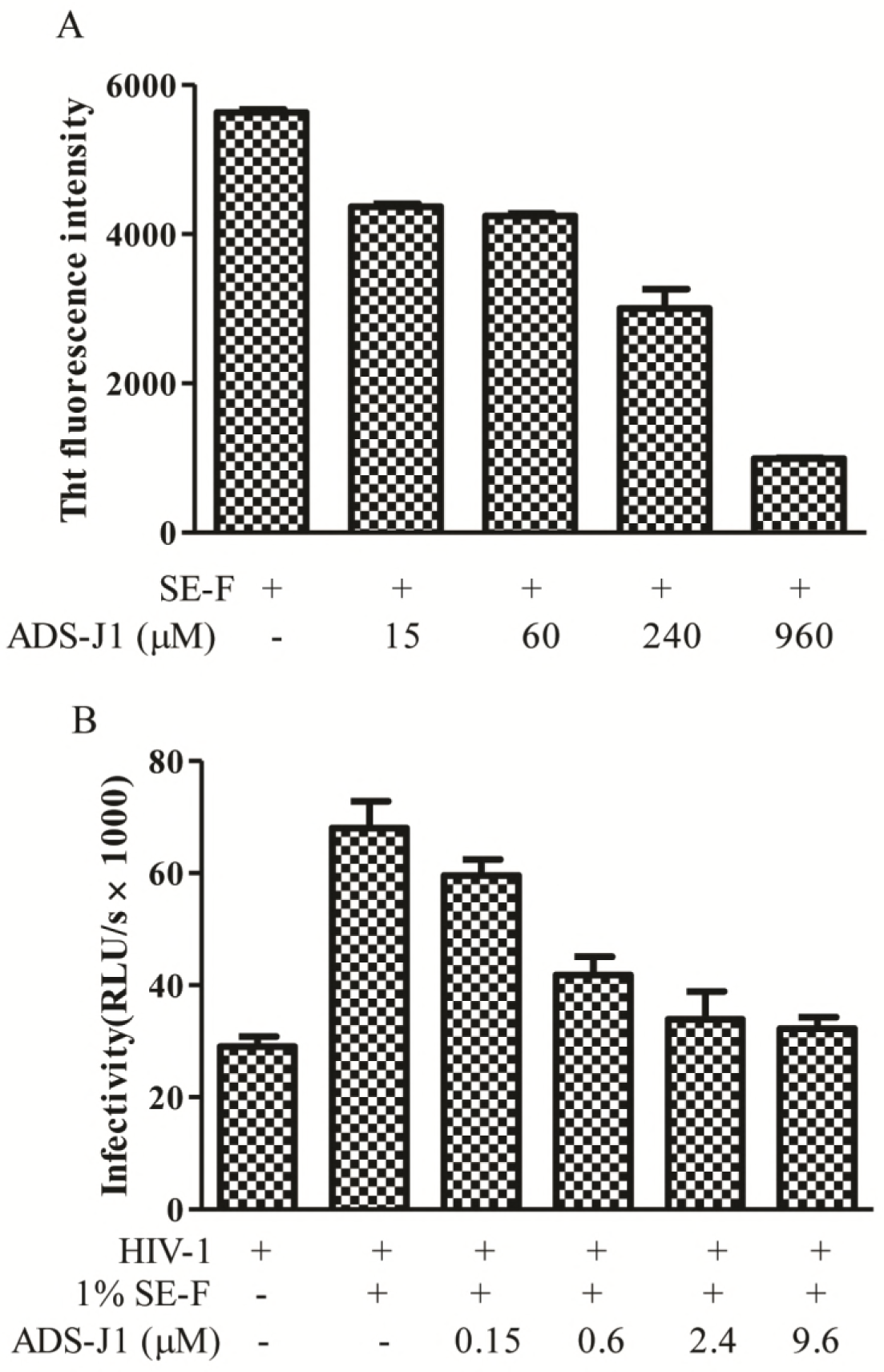
Disaggregating effects of ADS-J1 on endogenous seminal fibrils. (*A*) Loss of ThT fluorescence after incubation of SE-F with various concentrations of ADS-J1 under constant agitation was monitored at the time point of 8 h. Values represent mean fluorescence intensities derived from triplicate measurements ± SEM. (*B*) SE-F was exposed to ADS-J1 at different concentrations at 37°C for 8 h. Mixtures were then centrifuged and pellets were incubated with HIV-1 SF162. Subsequently, mixtures were used to infect TZM-bl cells. Luciferase activity were determined after 48 h. Results are means ± SEM from triplicate measurements.

### ADS-J1 converting SEVI fibrils into monomers

We proceeded to investigate the ADS-J1’s mode of action to remodel SEVI. Results from Tricine-SDS-PAGE (25, 26) revealed that the free supernatant of SEVI fibrils ran with a molecular weight consistent with a PAP_248-286_ monomer, which indicated that SEVI fibrils contained peptide monomer (Fig. 4 *A*). When treated with increasing level of ADS-J1, the supernatant of SEVI displayed increasing intensity of PAP_248-286_ monomer, compared with SEVI control. Quantification of bands showed higher proportion of monomer in samples treated with ADS-J1 (Fig. 4 *B*). Compared with EGCG, a universal amyloid fibril breaker, limited smear or trapped material was seen in SEVI with EGCG. EGCG is a well-known antioxidant chemical. In our experiments, after incubation of EGCG with SEVI, the solution changed appearance from transparent to brown, suggesting EGCG underwent auto-oxidation. In the presence of superoxide dismutase (SOD), a molecule that could decrease the auto-oxidation of EGCG (27), remodeling activities of EGCG were hampered by the rescue in ThT fluorescence compared with that in the absence of SOD. However, the remodeling effects of ADS-J1 were not influenced by SOD (Fig. 4 *C*).

**FIGURE 4.**
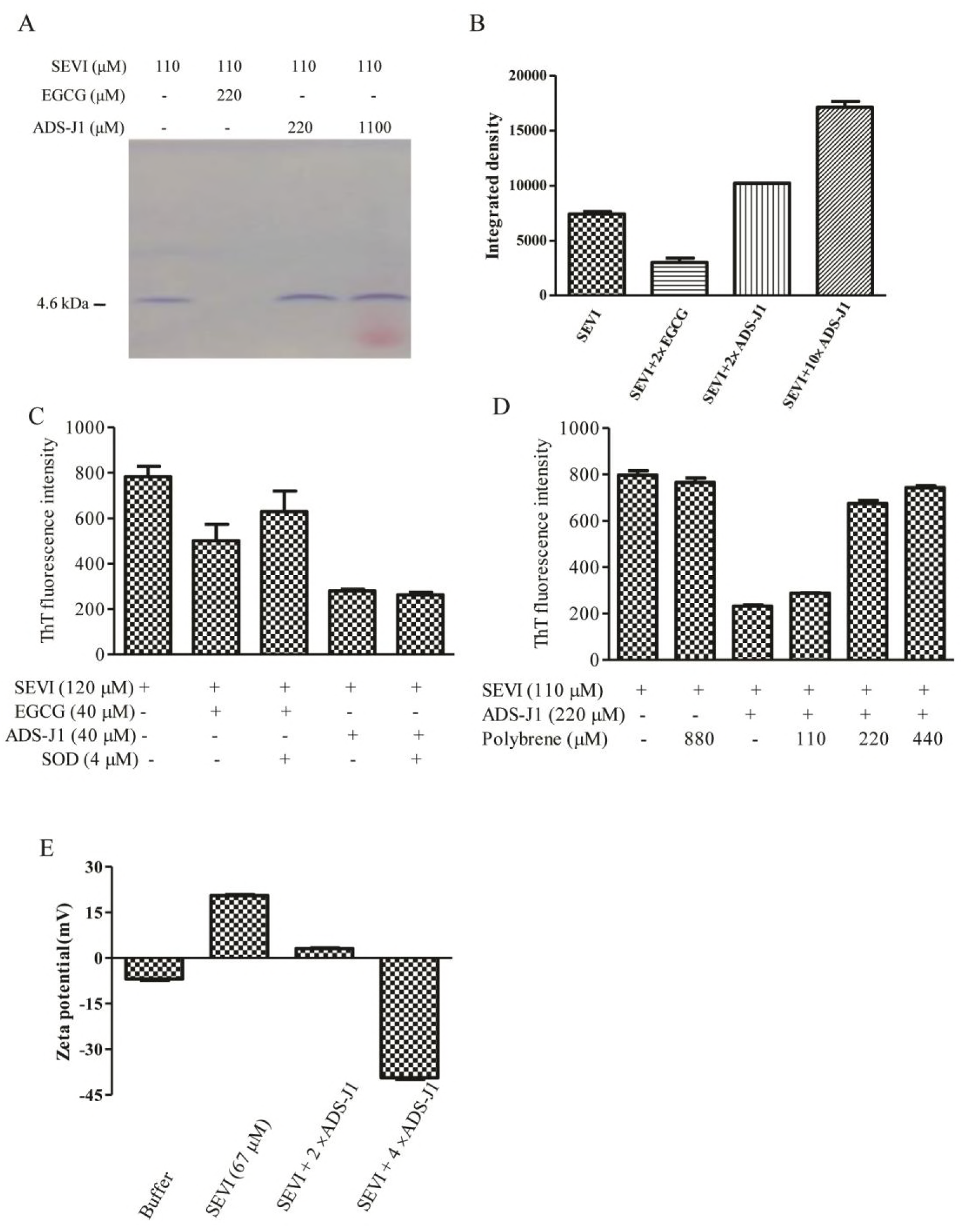
Disaggregation of SEVI fibrils into monomers by ADS-J1. (*A*) PAP_248-286_ monomer increased after SEVI was treated with ADS-J1. SEVI was agitated with ADS-J1 or EGCG and centrifuged at 12,000 rpm for 15 min. The supernatants were electrophoresed in Tricine-SDS-PAGE. Gel was stained with Coomassie blue. Positions of molecular weight standards are shown on the left. (*B*) Bar chart shows the quantification of peptide levels correspond to the main band in the Coomassie blue staining by using Quantity One image analysis software. (*C*) The remodeling activities of ADS-J1 were independent of auto-oxidation. SEVI was agitated with ADS-J1 or EGCG in the presence or absence of SOD for 4 h. Mixtures were centrifuged. The remaining fibrils in the pellets were determined by ThT staining assay. Values represent mean ThT fluorescence intensity derived from triplicate measurements ± SEM. (*D*) The effects of ADS-J1 on remodeling SEVI were counteracted by polybrene. ADS-J1 was incubated with different concentration of polybrene at 37°C for 30 min and centrifuged. The supernatants were agitated with SEVI and fibril integrity at the time point of 4 h was assessed by ThT staining assay. Results are means ± SEM of triplicate measurements.

Using a competitive experiment, we also found that pre-treatment of ADS-J1 with polybrene, a cationic polymer, reversed the fibril remodeling properties of ADS-J1 (Fig. 4 *D*). Polybrene displayed no effects on SEVI fibrils integrity. However, pre-treatment with increasing levels of polybrene, ADS-J1 gradually lost the ability to remodel SEVI fibrils. At an equimolar ratio of ADS-J1-polybrene (1:1), polybrene nearly reversed the remodeling effects of ADS-J1 on SEVI fibrils (Fig. 4 *D*), suggesting that polybrene and fibrils compete for binding to ADS-J1. The zeta potential measurement revealed that ADS-J1 neutralized the positive surface charge of SEVI. The zeta potential of SEVI (67 μM significantly decreased from 20.5±0.6 mv to −39.4±0.8 mv, as the concentration of ADS-J1 was increased from 0 μM to 268 μM (Fig. 4 *E*). These results show that ADS-J1 could disaggregate SEVI fibrils into monomer and electrostatic interaction, but not oxidation function, is a key determinant, differing from that of EGCG.

### Strong binding potency of ADS-J1 to PAP_248-286_ revealed by MD simulation

MD simulation was applied to investigate the potential mechanism of the remodeling activities of ADS-J1 (Fig. 5 *A*) on mature SEVI fibrils. Figure 5 *B* showed the structure of the electrostatic surface of PAP_248-286_ (positive charge shown in blue) with ADS-J1 molecules. PAP_248-286_ interacted with eight ADS-J1 molecules, which fully covered on the surface of PAP_248-286_. Since electrostatic interactions play vital roles in the remodeling effects of ADS-J1 on SEVI fibrils, we analyzed the ionic interactions between the S atom on ADS-J1 molecules and the N atom of the lysine side chain (or carbon atom of the guanidinium group of arginine) in PAP_248-286_. All eight positive charged residues in PAP_248-286_ were bound to ADS-J1 (Fig. 5 *C*). It showed the minimum X-S distances in the initial structure and average structure respectively for each residue.

**FIGURE 5.**
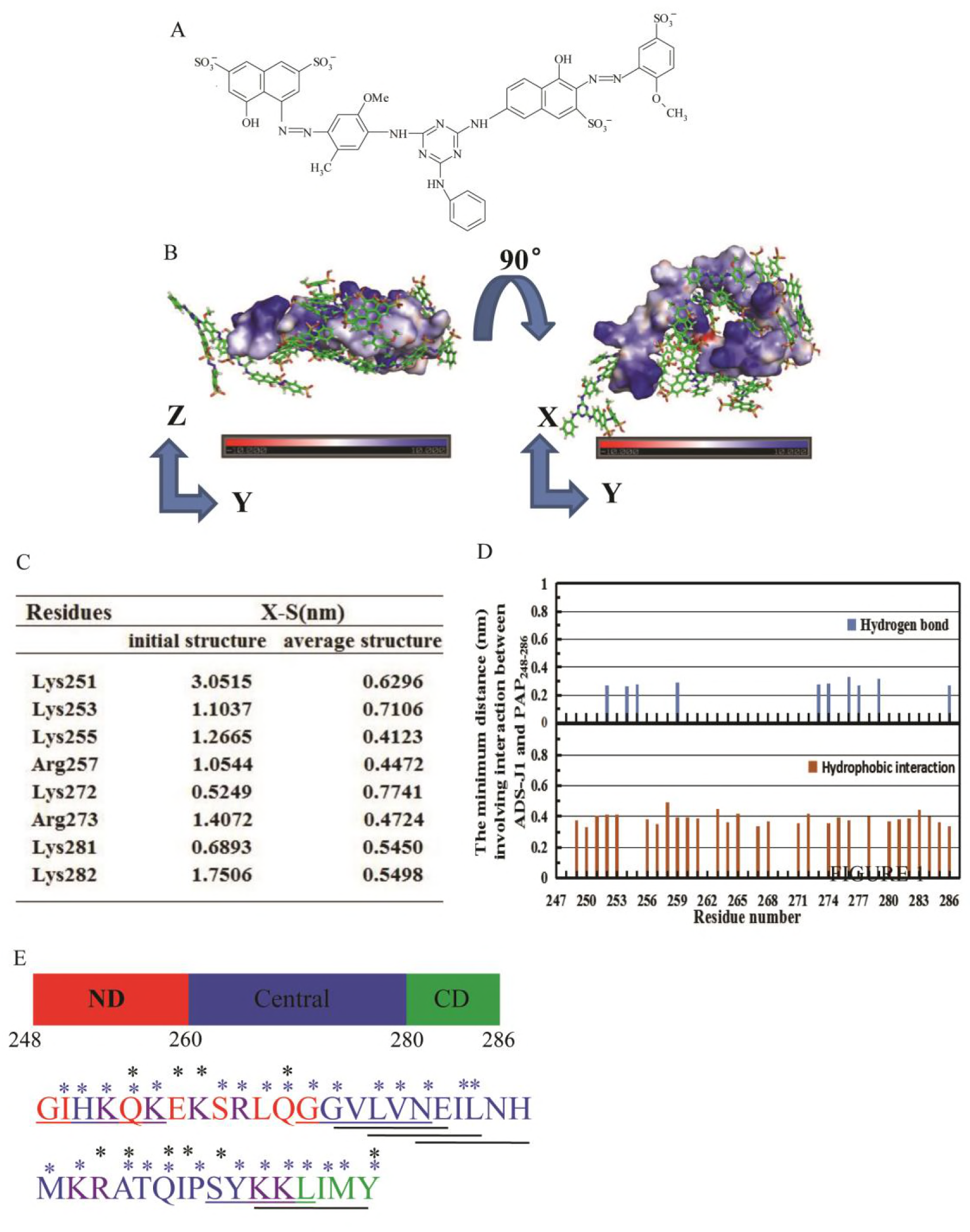
Representative PAP_248-286_-ADS-J1 binding complexes and potential interactions derived from MD simulation. (*A*) Chemical structure of ADS-J1. (*B*) The bottom (left panel) and back (right panel) views of the structure of PAP_248-286_ with eight ADS-J1 molecules. PAP_248-286_ is shown its electrostatic potential and colored according to the charge of atoms. Positively and negatively charged zones are visualized in blue and red, respectively. ADS-J1 is shown as stuck and colored by element. (*C*) ADS-J1 establishes electrostatic interaction with PAP_248-286_ as shown by decreased X-S distances (nm) between the S atom of sulfonic groups on ADS-J1 molecules and the N atom of the lysine side chain (or carbon atom of the guanidinium group of arginine) of the average structure, compared to that of the initial structure. (*D*) The minimum distance of ADS-J1 interacted with PAP_248-286_ residues via hydrogen bond and hydrophobic interaction. (*E*) Sequence of PAP_248-286_. The N-terminal domain (ND), central domain and C-terminal domain (CD) are depicted in red, blue and green respectively. The residues colored in purple indicate the positively charged amino acid. Black asterisks and blue asterisks represent the PAP_248-286_ residues interacting with ADS-J1 via hydrogen bond and hydrophobic interaction respectively. Predicted steric-zipper hexapeptides are underlined.

Besides electrostatic interactions, PAP_248-286_ and ADS-J1 also formed lots of hydrophobic interactions and hydrogen bonds (Fig. 5 *D*). Most of the PAP_248-286_ residues contacted with ADS-J1 molecules via hydrophobic interactions. These results suggested that almost all residues in PAP_248-286_ (35 out of 39) were occupied by ADS-J1 molecules (Fig. 5 *E*). Electrostatic interaction, hydrogen bond and hydrophobic interaction are the main driving forces for the binding of ADS-J1 to the PAP_248-286_.

### Secondary structure of PAP_248-286_ in the presence of ADS-J1 differing from that in the absence of ADS-J1

ADS-J1 binding results in important conformational changes in PAP_248-286_. The contact maps show the distances between pairs of amino acid residues in Figure 6 *A*. PAP_248-286_ alone adopts a kinked conformation, in which the C-terminus interacts with the N-terminus via hydrogen bonds leading to a U-turn in the central region. However, ADS-J1 binding converts PAP_248-286_ from the compact conformation to extended one. The average distances between the N-terminal Arg10/Leu11 with Ile37/Met38/Tyr39 in C-terminal were calculated over the two MD trajectories to further explain the difference between the conformational ensembles of PAP_248-286_ and PAP_248-286_ with ADS-J1 molecules. Figure 6 *B* and 6 *C* show that the distances between Arg10/Leu11 and Ile37/Met38/Tyr39 respectively keep stable around 0.5 nm in the absence of ADS-J1. In the presence of ADS-J1, however, the distances between Arg10-Tyr39, Leu11-Tyr39 and Arg10-Met38 become greater than 1 nm and distances between Arg10-Ile37, Leu11-Met38 and Leu10-Ile37 increase to more than 1.5 nm.

**FIGURE 6.**
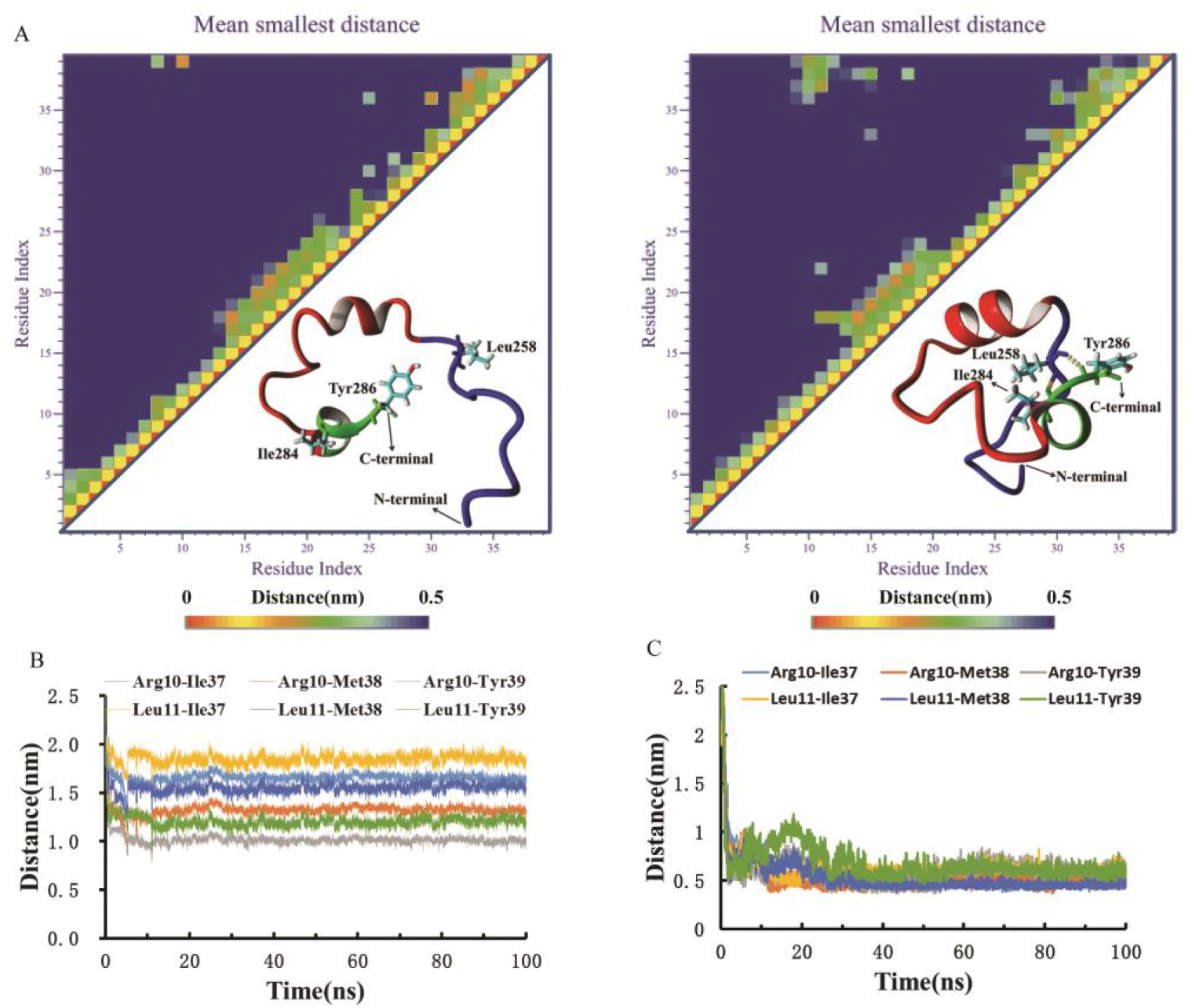
Induction of conformational changes of PAP_248-286_ by ADS-J1. (*A*) Contact map of residues in PAP_248-286_ in the presence (left) and absence of ADS-J1 (right). The inserts are the most representative structures of PAP_248-286_ obtained from the 100 ns simulation. ADS-J1 molecules are not shown. (*B-C*) The time-dependant changes of distance between the mass center of the Arg10-Ile37, Arg10-Met38, Arg10-Tyr39, Leu11-Ile37, Leu11-Tyr39 in the presence (*B*) and absence (*C*) of ADS-J1.

### Remodeling effects of ADS-J1 on a wide spectrum of amyloid fibrils

A number of amyloid fibrils are the pathogenic factors of several diseases, such as Aβ_1-42_ in Alzheimer’s disease and Amylin (fibrillar form of hIAPP_1-37_) in type Ⅱ diabetes (28, 29). In our previous study, we showed that EP2 derived from HIV-1 gp120 formed amyloid fibrils and greatly enhanced HIV-1 infection (30). We explored the remodeling effects of ADS-J1 on these amyloid fibrils. We found that ADS-J1 could effectively disassemble Aβ fibrils (Fig. 7 *A*, upper panel), amylin (Fig. 7 *B*, upper panel) and EP2 fibrils (Fig. 7 *C*, upper panel) in a dose-dependent manner, as evidenced by the decreased in ThT fluorescence after incubation of ADS-J1 with each fibril. TEM analysis was also applied to monitor the remodeling products. It is clear that the fibrils were reduced markedly and converted into some granular aggregates after incubation with ADS-J1 for 4 h, while untreated fibrils kept the well-developed fibrillar morphology (Fig. 7 *A*-*C*, bottom panel). The remodeling effects were also confirmed by particle size measurement. We observed that 4-fold excess of ADS-J1 drastically disrupted three fibrils into smaller structures (Fig. 7 *D*). These findings suggest that ADS-J1 could remodel a wide range of amyloid fibrils.

**FIGURE 7.**
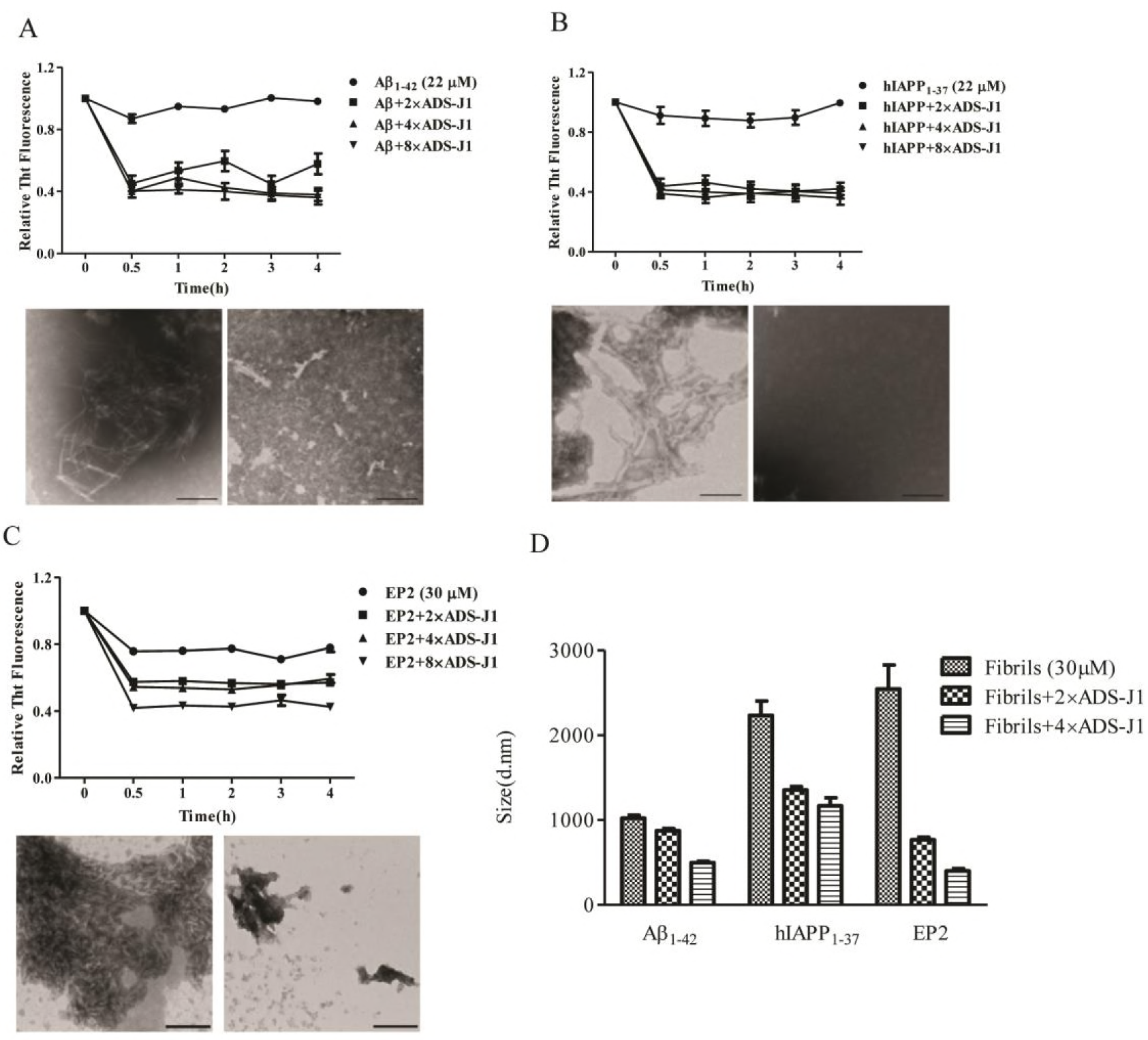
Remodeling effects of ADS-J1 on a wide spectrum of amyloid fibrils. ADS-J1 could remodel preformed fibrils of Aβ_1-42_(*A*), hIAPP_1-37_ (*B*) and EP2 (*C*) fibrils. Each of the mature fibrils were agitated in the presence or absence of ADS-J1 at molar ratio of fibril:ADS-J1=1:0,1:2,1:4,1:8. Aliquots were withdrawn at various time points and fibril integrities were assessed using ThT. Results are means ± SEM of three replicate experiments. The fibril structures in the absence (bottom, left) and presence (bottom, right) of ADS-J1 after agitation for 4 h with molar ratios of fibrils:ADS-J1 (1:4) were visualized by TEM. Scale bar = 100 nm. (*D*) Particle sizes of various fibrils agitated in the presence or absence of ADS-J1 for 4 h were measured. Results shown are means ± SEM of triplicate measurements from one of the two independent experiments, which yielded similar results.

## DISCUSSION

There is no effective cure to prevent HIV-1 sexual transmission topically. Therefore, therapeutic options capable to counteract the factors involved in the sexual transmission process are actively sought (12). In this study, we found that ADS-J1, apart from being an HIV-1 entry inhibitor, SEVI fibril inhibitor and blocker (3, 13, 31), was also an effective SEVI fibril disassembler. We found that ADS-J1 could effectively disrupt seminal amyloid fibrils, including SEVI fibrils and SEM1 fibrils as evidenced by loss of ThT fluorescence, disappearance of fibrils in TEM images, reduced particle size (Fig.1), impaired ability to enhance HIV-1 infection (Fig.2) and accumulated peptide monomer in remodeled products (Fig. 4, *A* and *B*). ADS-J1 could also disrupt endogenous amyloid fibrils in semen and impair its ability to enhance HIV-1 infection (Fig. 3). Importantly, our results showed that ADS-J1 could effectively disrupt SEVI and SEM1 fibrils within 4 hours, a time scale that would be critical for prevention of HIV-1 infection (Fig. 1).

ADS-J1 is also capable of disassembling multiple disease-related amyloid fibrils, such as Aβ_1-42_ fibrils, hIAPP_1-37_ fibrils, and EP2 fibrils (Fig. 7). As we known, there is limited effective amyloid reversing agent lies in the exceptional stability of the amyloid fibrils, which possess high resistance to proteases, protein denaturants, temperatures up to 98°C and 2% SDS (11). So far, the most reported fibril breakers can be divided into three categories (32). The first class is peptide based inhibitors, which are derived from different regions of the amyloidogenic peptide sequence. The homologous fragment could bind to a complementary sequence in the full-length protein, which results in interfering with subsequent protein self-assembly and favoring to disassemble preformed fibrils (33, 34). Some drawbacks have limited the clinical use of β-sheet breaker peptides, such as the cost of preparation, hydrosolubility, and oral bioavailability. The second class is ionic agents. Similar with ADS-J1, CLR01, a Lys-specific molecular tweezer with its functional phosphate groups, has been reported to be capable to disassemble several amyloid fibrils as those also investigated in this study (35–37). Of note, 2-fold excess of ADS-J1 could effectively remodel SEVI fibrils compared with CLR01 which required 10-fold excess (35). In addition, CLR01 fails to disassemble SEM1 fibrils and it takes several weeks to disassemble Aβ or α-synuclein (35), while ADS-J1 displays strong ability to remodel a wide range of amyloid fibrils within 4 h (Fig. 7).The third class of drugs is some well-known polyphenols, including EGCG, resveratrol, curcumin, oleuropein and scyllo-inositol etc.(32) Our results suggest that the disassembling process of ADS-J1 on SEVI fibrils is distinct from that of EGCG, a universal fibril breaker to date. The remodeling effects of EGCG on SEVI fibrils appear to be dependent on its auto-oxidation function (Fig. 4 *C*). EGCG undergoes auto-oxidation and generates superoxide and quinones in the presence of oxygen (38). PAP_248-286_ has six lysines and it is well established that quinones can react with the free amines comprising the N-terminus or the lysine side chains of proteins. It’s possible that oxidized EGCG induce profound chemical modification of the peptide side chains, resulting in disruption the β-sheet alignment and fibril disassembly. It could explain why the remodeling products of SEVI fibrils by EGCG were not recognized by SDS PAGE (Fig. 4 *A*). The antioxidant function or the auto-oxidation ability of EGCG might be responsible for its ability to interfere with a number of amyloid fibrils and to interact with numerous protein targets involved in various diseases, including cancer, metabolic, neurodegenerative, inflammation and microbial diseases (39). However, to be succeed in clinical application, several problems are still to be solved including, solubility, stability, potential hepatotoxicity (40, 41), inadequate target specificity and potential antagonizing effects in combination therapy (42, 43). ADS-J1 exerts limited oxidative capability. It might be associated with less host cytotoxicity and adverse events.

We then questioned why ADS-J1 poses excellent ability to disassemble various amyloid fibrils. We used SEVI as a model protein to elucidate the structure details of ADS-J1’s binding mode. Its structure might be a key determinant since it has not only ion group and also benzene and naphthalene as hydrophobic sidewall (Fig. 5 *A*). ADS-J1 displays strong binding propensity to full-length PAP_248-286_ via not only electrostatic interaction but also hydrophobic interaction and hydrogen bond. These results were similar with the observation that CLR03, a derivative of CLR01, lost the ability to disassemble amyloid fibrils because of lacking hydrophobic side chains (35). Based on these findings, we proposed a possible mechanistic model to interpret the disassembling effects of ADS-J1 on SEVI fibrils in Figure 8. ADS-J1 might firstly attach to the surface of SEVI fibrils, possibly via the positively charged residues, which are essential for their enhancement of HIV-1 infection. Then the strong binding propensity, including hydrogen bonds and hydrophobic interactions, leads to the penetration of ADS-J1 into the inner fibrillar structure by occupying the amyloid core sequence. ADS-J1 engaged key residues, which reside in predicted steric zippers (Fig. 5 *E*) (11, 44). Invaded by ADS-J1 molecules, fibrillar structures might be gradually loosened and cut into oligomers and then eventually monomers (Fig. 8 *B*).

**FIGURE 8.**
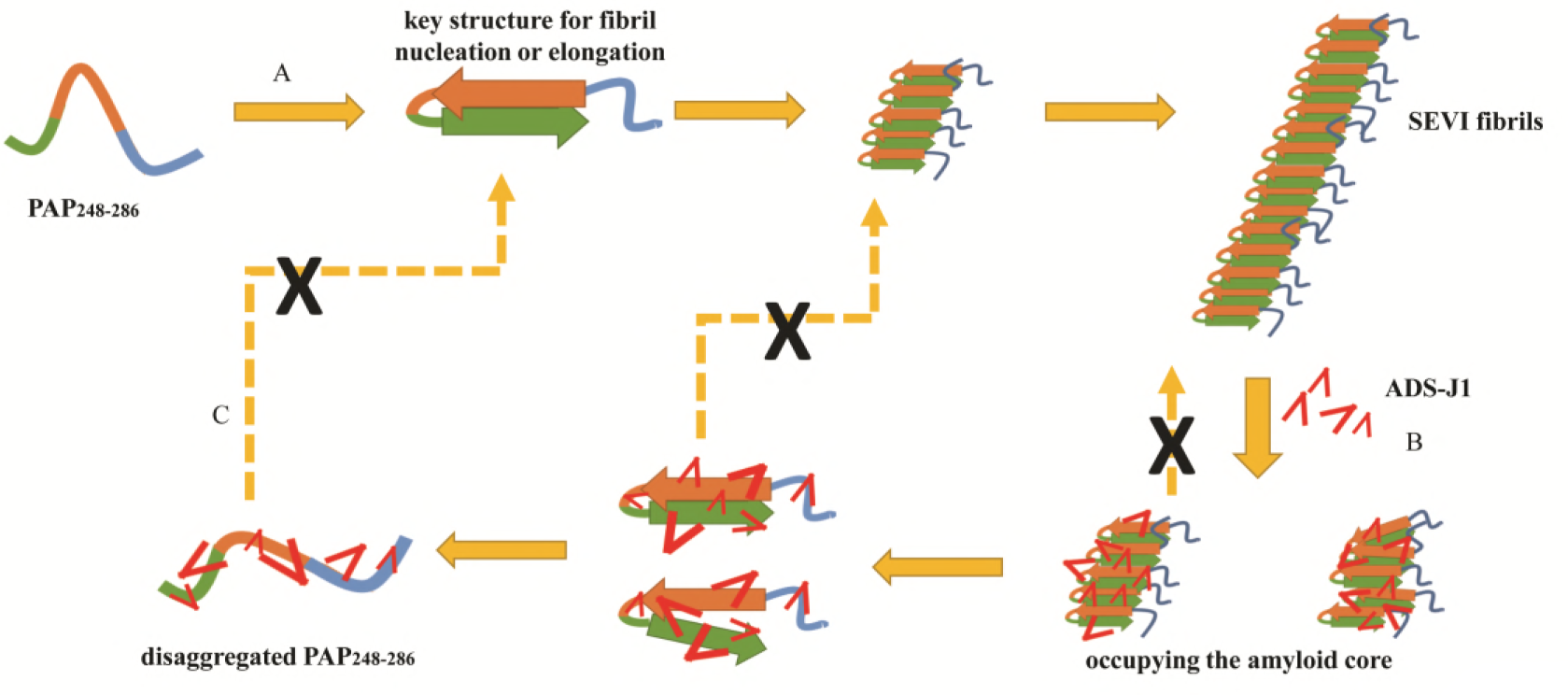
Schematic representation for the potential mechanism by which ADS-J1 disassembling SEVI fibrils. (*A*) PAP_248-286_ adopts a compact conformation in which the N-terminus interacts with C-terminus, leaving a deep U-turn in the central region. This conformation might be critical for the fibril nucleation and elongation. (*B*) ADS-J1 might firstly bind to the fibrils partially via electrostatic interaction. Then ADS-J1 molecules penetrate into the fibrils, occupying the amyloid core regions via hydrogen bonds, hydrophobic interactions and ionic interactions. Lastly, fibrils were degraded into oligomers and eventually monomers. (*C*) The disaggregated PAP_248-286_ might lose the vital conformation resulting in its deficiency in re-aggregating into fibrils.

Furthermore, maintaining specific conformational states is reported to be vital to form beta-sheet rich amyloid fibrils (45). As revealed by the MD images, PAP_248-286_ alone might adopt a compact conformation, in which the C-terminus interacts with the N-terminus leading to a deep U-turn in the central region. This unique structure might be essential to facilitate peptide aggregation (Fig. 8 *A*). However, ADS-J1 hinders the interactions between the C-terminal and the N-terminus and disturbs this potential aggregation-prone structure. Therefore, ADS-J1 could completely eliminate SEVI fibrils and yield fragments with reduced ability to form fibrils again due to the disfavoring conformations in the monomer capable of nucleation or elongation (Fig. 8 *C*).

Collectively, our study shows that ADS-J1 is broad-spectrum of amyloid fibrils breaker. It might be a suitable option for use as a topical microbicide component since it is an effective inhibitor and breaker of seminal fibrils originating from various amyloidogenic peptides. Anti-amyloid treatment will synergize with antiviral therapy, which appears encouraging for assessing therapies against the multifactorial nature of HIV-1 sexual transmission. In addition, the fact that ADS-J1 disaggregates various amyloid fibrils opens new perspectives for pharmaceutical applications and stimulates the synthesis of new, patentable agents towards amyloidogenic diseases.

## AUTHOR CONTRIBUTIONS

L. Q., Y. L., W. L. and L. H. performed and analyzed the experiments. Z. Y., H.L. and T. Z. performed MD experiments. T. Q. and Y. Q. collected the semen samples. L. L. and X. Z. discussed and modified the manuscript. L. Q., S. T. and S. L. co-wrote the paper. S. T. and S. L. designed and mastered the project process. The authors declare that they have no conflicts of interest with the contents of this article.

